# Fronto-limbic dysconnectivity leads to impaired brain network controllability in young people with bipolar disorder and those at high genetic risk

**DOI:** 10.1101/222216

**Authors:** Jayson Jeganathan, Alistair Perry, Danielle S. Bassett, Gloria Roberts, Philip B. Mitchell, Michael Breakspear

## Abstract

Recent investigations have used diffusion-weighted imaging to reveal disturbances in the neurocircuitry that underlie cognitive-emotional control in bipolar disorder (BD) and in unaffected siblings or children at high genetic risk (HR). It has been difficult to quantify the mechanism by which structural changes disrupt the superimposed brain dynamics, leading to the emotional lability that is characteristic of BD. Average controllability is a concept from network control theory that extends structural connectivity data to estimate the manner in which local neuronal fluctuations spread from a node or subnetwork to alter the state of the rest of the brain. We used this theory to ask whether structural connectivity deficits previously observed in HR (*n*=84, mean age 22.4) individuals, patients with BD (*n*=38, mean age 23.9), and age- and gender-matched controls (*n*=96, mean age 22.6) translate to differences in the ability of brain systems to be manipulated between states. Localized impairments in network controllability were seen in the left parahippocampal, left middle occipital, left superior frontal, right inferior frontal, and right precentral gyri in BD and HR groups. Subjects with BD had distributed deficits in a subnetwork containing the left superior and inferior frontal gyri, postcentral gyrus, and insula (*p*=0.004). HR participants had controllability deficits in a right-lateralized subnetwork involving connections between the dorsomedial and ventrolateral prefrontal cortex, the superior temporal pole, putamen, and caudate nucleus (*p*=0.008). Between-group controllability differences were attenuated after removal of topological factors by network randomization. Some previously reported differences in network connectivity were not associated with controllability-differences, likely reflecting the contribution of more complex brain network properties. These analyses highlight the potential functional consequences of altered brain networks in BD, and may guide future clinical interventions.

**Highlights:**

- Control theory estimates how neuronal fluctuations spread from local networks.
- We compare brain controllability in bipolar disorder and their high-risk relatives.
- These groups have impaired controllability in networks supporting cognitive and emotional control.
- Weaker connectivity as well as topological alterations contribute to these changes.

## 1. Introduction

For much of the 20^th^ century, neuroscience was predicated on the notion that individual cognitive functions could be attributed to segregated regions of the brain. In recent times, this paradigm has shifted toward a connectivity-based approach which emphasizes the crucial role of network-mediated functional integration (Sporns, 2013). In parallel, the clinical neurosciences have shifted from a predominantly lesion-based approach towards a connectomic framework (Fornito *et al.*, 2015). However, disrupted network connections in the brain impact not only directly connected regions, but can also influence distant cortical regions through complex dynamics (Crofts *et al.*, 2011). To characterize the spread of dysfunction in disease states, network connectivity and the superimposed dynamics of brain activity must both be considered (Stam, 2014). In human brains, the former is now readily accessible through advances in diffusion-weighted imaging and tractography (Jbabdi *et al.*, 2015; Farquharson and Tournier, 2016). However, the principles of neuronal dynamics, well-known at the neuron level, remain incompletely understood at the macroscopic scale (Breakspear, 2017).

Studies using diffusion-weighted imaging have identified ‘disconnection syndromes’ in schizophrenia (Stephan *et al.*, 2009; Zalesky *et al.*, 2011), depression (Bai *et al.*, 2012), attention-deficit hyperactivity disorder (Cao *et al.*, 2013), and epilepsy (Widjaja *et al.*, 2015). More recently, attention has turned to bipolar disorder (BD), a psychiatric condition characterized by episodic disturbances in mood and cognition. While variations in mood around the set point of euthymia are intrinsic to human experience, excursions of mood in BD reach a magnitude and duration that produces substantial distress, dysfunction, and disability. Recent structural imaging studies of BD have consistently reported white matter alterations in circuits involving prefrontal, striatal, and limbic regions (Xekardaki *et al.*, 2011; Nortje *et al.*, 2013; Phillips and Swartz, 2014; Roberts *et al.*, 2016b). The disease also has a strong genetic component. Unaffected first-degree relatives often have attenuated affective disturbances and are at significantly increased risk of developing the disorder; their odds-ratio is estimated to lie between 7 and 14. These relatives are hence considered at high risk (HR), particularly younger relatives who have not passed the peak age of onset (Weissman *et al.*, 1986; McGuffin *et al.*, 2003; Mortensen *et al.*, 2003; Perlis *et al.*, 2004; Purcell *et al.*, 2009; Perich *et al.*, 2015). Identification of structural and functional connectivity differences between BD patients, their unaffected relatives, and healthy controls, have permitted a dissociation of genetic, illness-expression, and adaptive influences (Frangou, 2009; Kempton *et al.*, 2009; Pompei *et al.*, 2011; Sprooten *et al.*, 2011; Meda *et al.*, 2012; Doucet *et al.*, 2017; Ganzola *et al.*, 2017). The neurodevelopmental risk of their first-degree relatives is supported by effective and functional connectivity studies that reveal alterations in fronto-limbic networks supporting emotion processing and regulation (Pompei *et al.*, 2011; Frangou, 2012; Breakspear *et al.*, 2015; Dima *et al.*, 2016). We recently investigated structural network disturbances in both BD patients and youth at high genetic risk for the disorder (Roberts *et al.*, 2016b). The high-risk (HR) group demonstrated unique disturbances in subnetworks of connections centering upon the inferior frontal gyri and insular cortex. These findings speak to an endogenous risk, present as these subjects transition through a critical developmental period.

Structural connectivity deficits on their own, without an informed understanding of network topology or the overlying dynamics, comprise ‘correlation’ rather than ‘causation’. Relating structural network disturbances in psychiatric conditions to the illness phenotype is an important yet unfulfilled ambition. Whole-brain simulations of large-scale neuronal dynamics could theoretically achieve this goal, but their role is challenged by the complex nature of spatially heterogeneous local dynamics and limitations in current techniques for inferring nonlinear dynamics from noisy imaging data. Here, we tackle this issue with network control theory, which combines structural connectivity with a linear spatially homogenous estimate of local dynamics to model and predict the spread of neuronal signals within the brain. The extent of signal spread is quantified by controllability. Note that this analysis rests exclusively upon the mathematical analysis of structural connectivity data. It yields *predicted* changes in brain dynamics under the assumption that stable (linear) approximations serve as a useful guide to actual brain dynamics and, by inference, corresponding cognitive processes.

Network control theory has recently been used to predict brain dynamics and stability, with the understanding that brain regions whose structural connectivity suggest poor controllability over the rest of the brain, will indeed have poorer control over brain states (Gu *et al.*, 2015; Betzel *et al.*, 2016). As neuronal activity evolves in time, functional brain states undergo transitions, traversing a path through a dynamic state-space landscape. Perturbations applied to a set of control nodes, either from an extrinsic source or from internal dynamics, can modulate these trajectories (Bassett and Khambhati, 2017). This input energy depends on the choice of control nodes, and the strength and pattern of structural connections (Kim *et al.*, 2018; Wu-Yan *et al.*, 2018). Recent work has found intriguing links between the average controllability of different brain systems and performance during cognitive testing (Betzel *et al.*, 2016; Muldoon *et al.*, 2016; Tang *et al.*, 2017).

In general, high brain network controllability appears to reside in those cortical regions typically associated with executive function, self-monitoring and emotional control including the default, limbic and cognitive control systems (Gu *et al.*, 2015). The notion that dysregulation of affect in BD could arise from the structural disturbances impacting upon the controllability of these systems has intuitive appeal. We sought to test that hypothesis here. In other words, we hypothesize that brain regions whose structural connectivity suggests a lack of control over the rest of the brain, actually contribute to the poor emotional control seen in BD and in muted form, in their first-degree relatives. We first characterize the network underpinnings of average controllability, and then apply controllability analyses to brain regions and subnetworks where changes in bipolar and high-risk cohorts were previously reported (Roberts *et al.*, 2016b). We hypothesized that the structural dysconnectivity of these subnetworks and regions would be associated with impairments in wider network controllability, particularly in the neurocircuitry that underlies cognitive-emotional control.

## 2. Materials and methods

### 2.1. Participants

218 participants between 15-30 years of age comprise three richly-phenotyped groups: (i) 84 participants at high risk (HR) of bipolar disorder (mean age 22.4), (ii) 96 controls (CNs) without a family history of mental illness (mean age 22.6), and (iii) 38 bipolar disorder (BD) patients (mean age 23.9; 18 bipolar I, 20 bipolar II) (Roberts *et al.*, 2016b). HR and BD participants were recruited from families who had previously participated in a BD pedigree molecular genetics study or a specialized BD research clinic, as well as from clinicians, mental health consumer organisations, and other forms of publicity. Inclusion in the HR group was determined by having a first-degree relative with BD, with 19 in the HR group having a sibling with BD and 65 having a parent with BD. There were five families in this study that included both a participant in the HR group and the BD group. CN subjects were recruited via advertisements in print, electronic notices, and noticeboards in universities and local communities. Demographic and clinical data are provided in Table 1. Further description of the sample ascertainment and clinical assessments for the current population are provided in (Perich *et al.*, 2015).

**Table 1.** Demographic and clinical data.

|  | **Controls** | **High-risk** | **Bipolar disorder** | ***p*-value** | **Post-hoc Effects** |
| --- | --- | --- | --- | --- | --- |
| Number of subjects | 96 | 84 | 38 | - |  |
| Males, *n* (%) | 43 (44.8) | 39 (46.4) | 15 (39.5) | 0.77 |  |
| Intelligence Quotient, mean (SD) | 117.7 (10.3) | 116.3 (10.7) | 117.3 (12.0) | 0.69 |  |
| Age, mean (SD) | 22.6 (3.8) | 22.4 (4.7) | 23.9 (3.4) | 0.13 |  |
| **Lifetime DSM-IV diagnosis** |  |  |  |  |  |
| Any diagnosis, *n* (%) | 24 (25.0) | 39 (46.4) | 38 (100.0) | <0.001 | HR > CN**<br>BD > CN*** |
| At least one MDE, *n* (%) | 9 (9.4) | 22 (26.2) | 36 (94.7) | <0.001 | HR > CN**<br>BD > CN***<br>BD > HR*** |
| Recurrent MDD, *n* (%) | 1 (1.0) | 7 (8.3) | - | 0.01 | HR > CN* |
| Any anxiety disorder, *n* (%) | 9 (9.5) | 15 (18.3) | 15 (39.5) | <0.001 | BD > CN***<br>BD > HR* |
| Any behavioural disorder, *n* (%) | 1 (1.1) | 6 (7.4) | 7 (18.9) | <0.001 | HR > CN*<br>BD > CN*** |
| Any substance disorder, *n* (%) | 6 (6.3) | 9 (10.7) | 6 (15.8) | 0.220 | - |
| **Symptom severity scales** |  |  |  |  |  |
| 22 to 30 years | *n*=51 | *n*=41 | *n*=25 |  |  |
| MADRS, mean (SD) | 1.9 (3.2) | 2.5 (3.7) | 10.1 (9.5) | <0.001 | BD > CN*** |
|  |  |  |  |  | BD > HR*** |
| 15 to 21 Years | *n*=34 | *n*=33 | *n*=9 |  |  |
| CDI, mean (SD) | 6.8 (3.7) | 9.0 (6.6) | 21.7 (8.7) | <0.001 | BD > CN*** |
|  |  |  |  |  | BD > HR*** |
| **Age at First** |  |  |  |  |  |
| MDE, mean (SD) | 19.1 (3.2) | 18.5 (4.5) | 15.4 (3.7) | 0.002 | BD < CN** |
|  |  |  |  |  | BD < HR** |
| Manic episode, mean (SD) | - | - | 17.5 (3.2) | - | - |
| Mood episode, mean (SD) | 19.1 (3.2) | 18.5 (4.5) | 14.9 (3.8) | <0.001 | BD < CN** |
|  |  |  |  |  | BD < HR** |
| Any anxiety disorder | 10.3 (6.7) | 13.5 (6.4) | 13.3 (7.0) | 0.277 |  |
| **Psychotropic Medication** |  |  |  |  |  |
| Anti-depressants, *n* (%) | 0 (0.0) | 0 (0.0) | 15 (39.5) | - | - |
| Mood stabilisers, *n* (%) | 0 (0.0) | 0 (0.0) | 26 (68.4) | - | - |
| Anti-psychotics, *n* (%) | 0 (0.0) | 0 (0.0) | 11 (28.9) | - | - |

Significance Levels: \*\*\* $p < 0.001$ , \*\* $p < 0.01$ , \* $p < 0.05$ .

The study was conducted with approval from the University of New South Wales Human Research Ethics Committee (HREC Protocol 09/097) and the South Eastern Sydney Illawarra Health Service HREC (Protocol 09/104). Written informed consent was obtained from all participants.

### 2.2. Construction of structural networks

Diffusion-weighted images were obtained using a 3T Philips Achieva X MRI scanner and analyzed as previously described (Roberts *et al.*, 2016b). In short, pre-processed data underwent spherical deconvolution followed by probabilistic tractography to generate 5 million streamlines. The AAL template (Tzourio-Mazoyer *et al.*, 2002) was subdivided into 512 regions of uniform volume (Zalesky *et al.*, 2010b) (see https://github.com/AlistairPerry/CCA). Weighted structural networks were produced by combining each subject’s tractography with their parcellation template (Roberts *et al.*, 2016b). Edge weights represent the number of streamlines connecting two parcels. Structural networks were thresholded to maintain the 10% top-weighted connections, as investigations typically analyze brain networks with densities centering upon this value (Sporns, 2013; Perry *et al.*, 2015). Analyses of 5%, 15%, and maximum densities are provided in Supplementary Table 3, 4 and 5.

### 2.3. Network control theory

Linear control theory quantifies transient responses to brief perturbations of a linearly stable system. In this study, structural networks were overlaid with linear dynamics evolving in discrete time steps (*t*=1,2, …) according to, 

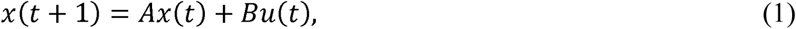

where *x*(*t*) is a vector denoting the simulated state of all nodes *i* at time *t*. The matrix *A* is the structural connectivity matrix, with element *A*_*ij*_ representing the number of white matter streamlines connecting regions (or *nodes*) *i* and *j*. The input matrix *B* specifies the control nodes in the brain (those receiving the perturbation), 

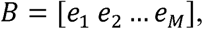

where *e*_*i*_ is a column vector with a ‘1’ in the position(s) corresponding to the control node(s). The variable *u*(*t*) is the energy applied to the set of control nodes *B* at time *t*.

The simulated states *x* represent underlying neural states (firing rates, membrane conductances, etc.), which in turn support the cognitive, perceptual, and emotional processes arising in the corresponding brain networks. Individual subject networks were normalized to prevent unstable dynamics (see Supplementary material).

Network control theory thus predicts the response of a complex system to perturbations, such as sensory (or physiological) perturbations. This linear system is a simplification of the full nonlinear neural dynamics that would more realistically embody the behavior of large-scale brain dynamics (Breakspear, 2017). However, linear controllability of a structural network is usually sufficient to imply controllability of nonlinear dynamics overlaid on the same structure, because linear dynamics are accurate around stable fixed points and periodic attractors (local, stable equilibria and oscillations) (Deco *et al.*, 2008). Our choice of discrete-time dynamics is based on prior work showing that controllability is similar in discrete- and continuous-time settings (Gu *et al.*, 2015). Note that the time steps of the model are arbitrary in the sense that shorter (or longer) time steps can be accommodated by simply scaling the magnitude of the elements of *A* and *B* up (or down).

Average controllability for a set of control nodes is the average energy needed to steer the system to any target state in finite time (Fig. 1). We use the mean impulse response energy as a measure of average controllability as it satisfies a relation of inverse proportionality. The impulse response is the total magnitude and extent of signal spread when a stimulus signal is applied to a node or subnetwork (see Supplementary material for further details). In the rest of this paper, we refer to average controllability simply as ‘controllability’ for brevity, noting that there are other independent controllability metrics that have been studied in the setting of structural brain networks (Wu-Yan *et al.*, 2018). The code pertaining to the calculation of average controllability from each participant’s structural network matrix is available for download from here (https://www.danisbassett.com/uploads/1/1/8/5/11852336/ave_control-2.m).

**Figure 1.**
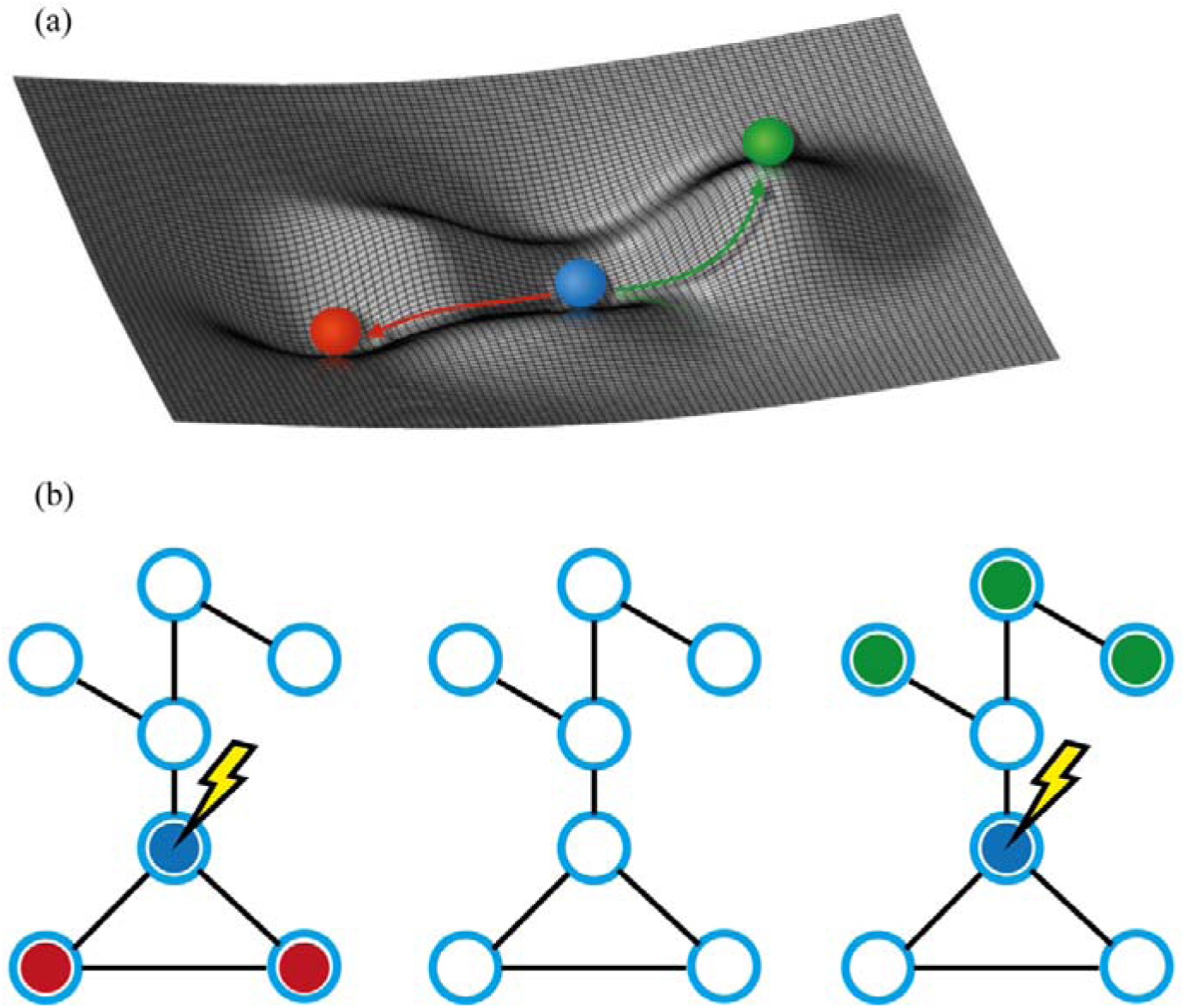
Average controllability. (a) Schematic representation of the mean impulse response energy required to steer a dynamic network from its base “resting” state (blue ball) to either an easy-to-reach state (red) or a hard-to-reach state (green). (b) Schematic representation of corresponding changes from the baseline state (centre) to an easy-to-reach state (left, red) or a hard-to-reach state (right, green). Arrow depicts the node receiving the perturbation.

### 2.4. Contributors to average controllability

Nodal strength is the weighted sum of each node’s connections to its neighbors on the network, and has an important influence on controllability (Gu *et al.*, 2015). However, other network features also influence controllability, such as the distribution of paths of different lengths as measured by the communicability matrix (Betzel *et al.*, 2016; Gu *et al.*, 2017) and the diversity of connections strengths from the controlled node(s) to the target node(s) (Kim *et al.*, 2018). Some topological factors, such as scale-freeness, are captured in the distribution of node strengths, while others, such as clustering, can occur independently of the strength distribution. How much variance each of these statistics accounts for in the controllability estimates can depend on the architecture of the specific graph under study (Wu-Yan *et al.*, 2018). Therefore, we first sought to disambiguate the contribution to network controllability of edge strength from higher order measures of network topology in our data.

To examine the impact of strength distribution on variability in controllability for each node in structural brain graphs, that node’s controllability was calculated before and after strength-preserving randomization (Maslov and Sneppen, 2002). If these values are perfectly correlated, then strength accounts for all of that node’s controllability: Lower correlations imply that other network features influence that node’s controllability. Topological network features can contribute to between-subjects and between-nodes variability in controllability. We assessed the presence and strength of such relations separately by correlating controllability values across subjects or across nodes (see Supplementary material for more detail). The coefficient of determination was calculated as the square of the Spearman rank correlation coefficient.

We next examined how other widely adopted nodal measures of network topology contribute to average controllability; this is important because our intuitions about brain graphs are heavily influenced by prior studies using these measures of network topology, independent of posited dynamics. To identify which of these measures are correlated with estimates of controllability, stepwise linear regression was implemented using entry and removal probabilities of 0.05 and 0.10, respectively. The nodal measures that were entered in the regression model were node strength, local clustering coefficient, subgraph centrality, local efficiency, betweenness centrality, within-module degree *z*-score, and participation coefficient. The latter two measures were calculated based upon the modular decomposition of the group average network from the healthy control participants, detected with a Louvain-like locally greedy algorithm for maximization of the modularity quality (Rubinov and Sporns, 2010). The nodal values for each node were averaged across all CN participants before the stepwise regression was performed.

### 2.5. Controllability at different levels of network granularity

We calculated group differences in average controllability metrics at three levels of network granularity: single nodes, subnetworks of nodes, and communities derived from intrinsic functional connectivity patterns.

#### 2.5.1. Node level

Our previous analyses of structural connectivity identified seven brain regions with differences in nodal strength between CN, HR, and BD groups (Roberts *et al.*, 2016b). Here, we calculated the average controllability of these seven nodes; that is, the ability to control whole brain dynamics through a stimulus to each of these individual nodes.

Group differences in average controllability were elicited with a one-way ANOVA for all seven nodes with a false discovery rate (FDR) correction. We then performed a one-tailed *t*-test of surviving nodes using previously identified group contrasts, and a final FDR correction step. The Benjamini-Hochberg procedure (α = 0.05) was used for FDR correction.

#### 2.5.2. Subnetwork level

We previously analyzed group effects in these data using the network-based statistic (NBS), a permutation-based method which employs topological inference to control for family-wise error when identifying subnetworks with group-wise differences in connection strength (Zalesky *et al.*, 2010a). Connectivity differences between CN, HR, and BD were observed in four subnetworks (Roberts *et al.*, 2016b) (Fig. 2). In this study, we calculated the controllability of all nodes within these subnetworks. Subnetwork average controllability is the sum of average controllability values for all nodes in the subnetwork. Intuitively, this measures the impulse response to a stimulus applied across the subnetwork. This impulse response depends on (i) the subnetwork’s external connectivity to surrounding regions, and (ii) the internal connections within the subnetwork, which amplify the signal by internal excitatory feedback (Fig. 3) (Kim *et al.*, 2018). Because these networks were discovered by the NBS, they have, by definition, weaker internal connectivity but relatively preserved external connectivity. Hence, subnetwork average controllability here is a surrogate for abnormal internal signal amplification.

**Figure 2.**
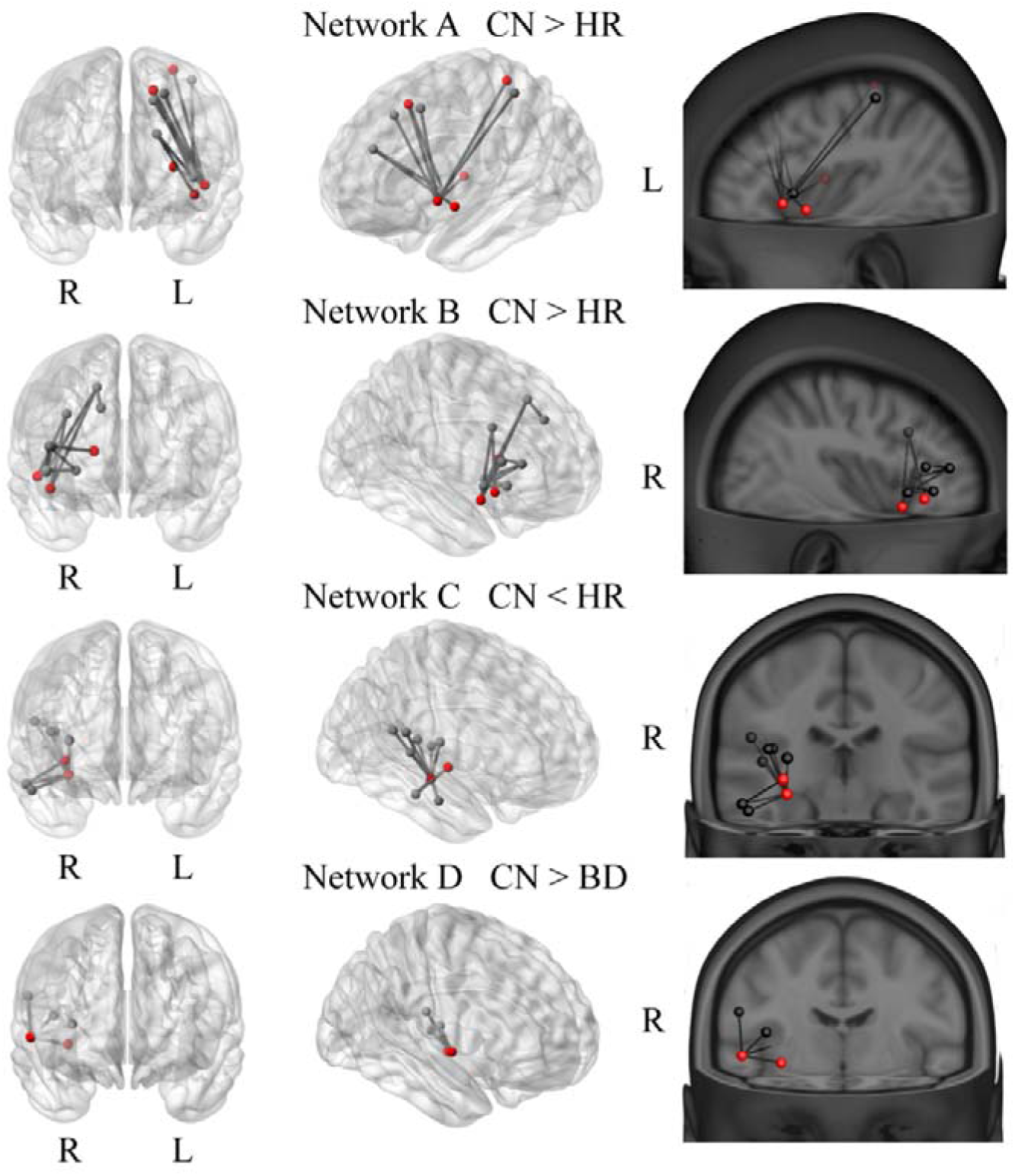
Subnetworks showing significant between-group differences as identified by the network-based statistic in the previous investigation. (Roberts *et al.*, 2016b). Connections (lines) between nodes (circles) with significant group differences in streamline count, with nodes coloured according to their previously identified node degree. Red circles indicate high-degree hub regions (top 15%), while grey circles represent non-hubs. CN, controls; HR, high risk; BD, bipolar disorder; L, left; R, right; α, azimuth. Figure adapted with permission from (Roberts *et al.*, 2016b).

**Figure 3.**
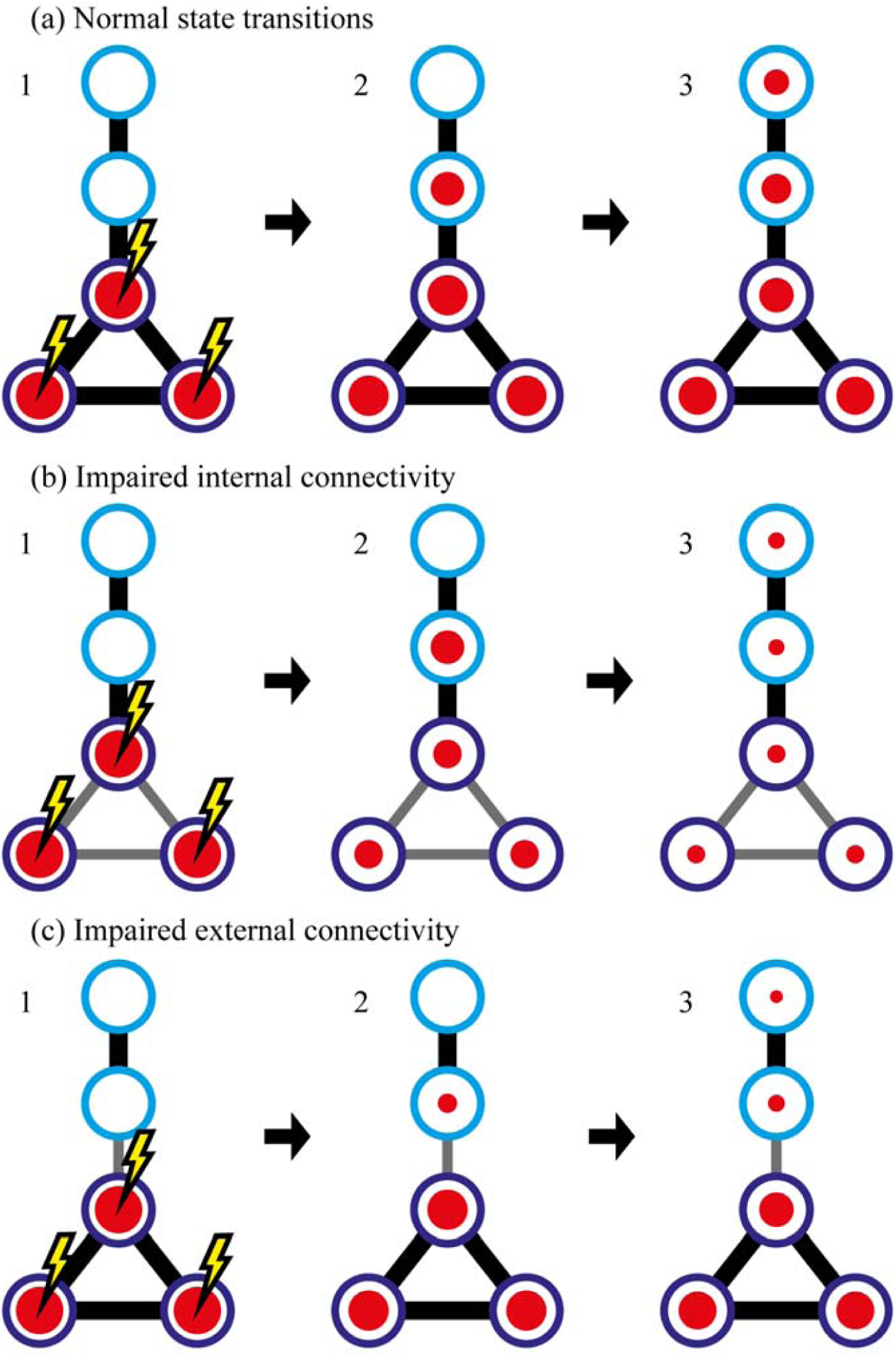
Schematic of signal spread from a closed loop of three nodes. (purple circles in left panels). The size of the circle indicates the corresponding magnitude of activation across the subnetwork. Attenuated intranetwork (b) or outgoing connections (c) (grey lines) can diminish the impulse response.

To ascertain the influence of network topology on group differences in subnetwork controllability, the subnetwork analysis was repeated after strength-preserving randomization. If the group differences were preserved, then these differences could be attributed primarily to differences in the strength of the nodes within each subnetwork. Changes in group differences following strength-preserving randomization reflect contributions from higher order topological features.

The following statistical protocol was used: a one-way ANOVA of group differences in each subnetwork’s average controllability, then FDR correction of the four resulting *p*-values. Two-tailed *t*-tests were then performed for surviving binary contrasts followed by a final FDR correction step for the individual possible group effects.

#### 2.5.3. Community level

Nodal-regions were assigned to widely adopted functional network affiliations derived from resting state fMRI data (Yeo *et al.*, 2011). The community affiliation of a brain-region was identified by its best spatial fit within one of the seven functional networks (Perry *et al.*, 2017) (see https://github.com/AlistairPerry/CCA). As with the subnetwork controllability, the average controllability of a community is the sum of the controllability values for all nodes within that community. A one-way ANOVA was first used to test for group-wise differences in the controllability of each community, followed by FDR-correction.

## 3. Results

### 3.1. Contributors to controllability

The nodal strength and average controllability of each node were positively correlated (*R*=0.89; *p*<0.0001, see Supplementary material for a formal derivation). This association is consistent with previous analyses of other structural connectomic data in the brain (Gu *et al.*, 2015) and non-brain graphical models (Wu-Yan *et al.*, 2018). Regions of high average controllability, with the potential to shift the rest of the brain easily between states, are topologically well-connected regions.

The brain network factors underlying average controllability were probed with a randomization scheme that preserved node-strength distribution within the network but destroyed higher-order topological properties. Strength distribution accounted for 87% of between-node variability in average controllability within a subject and 68% of between-subject variability. The remainder can hence be attributed to other topological factors. Figure 4 demonstrates the shrinkage of the mean and variance of subject-mean controllability values after network randomization.

**Figure 4.**
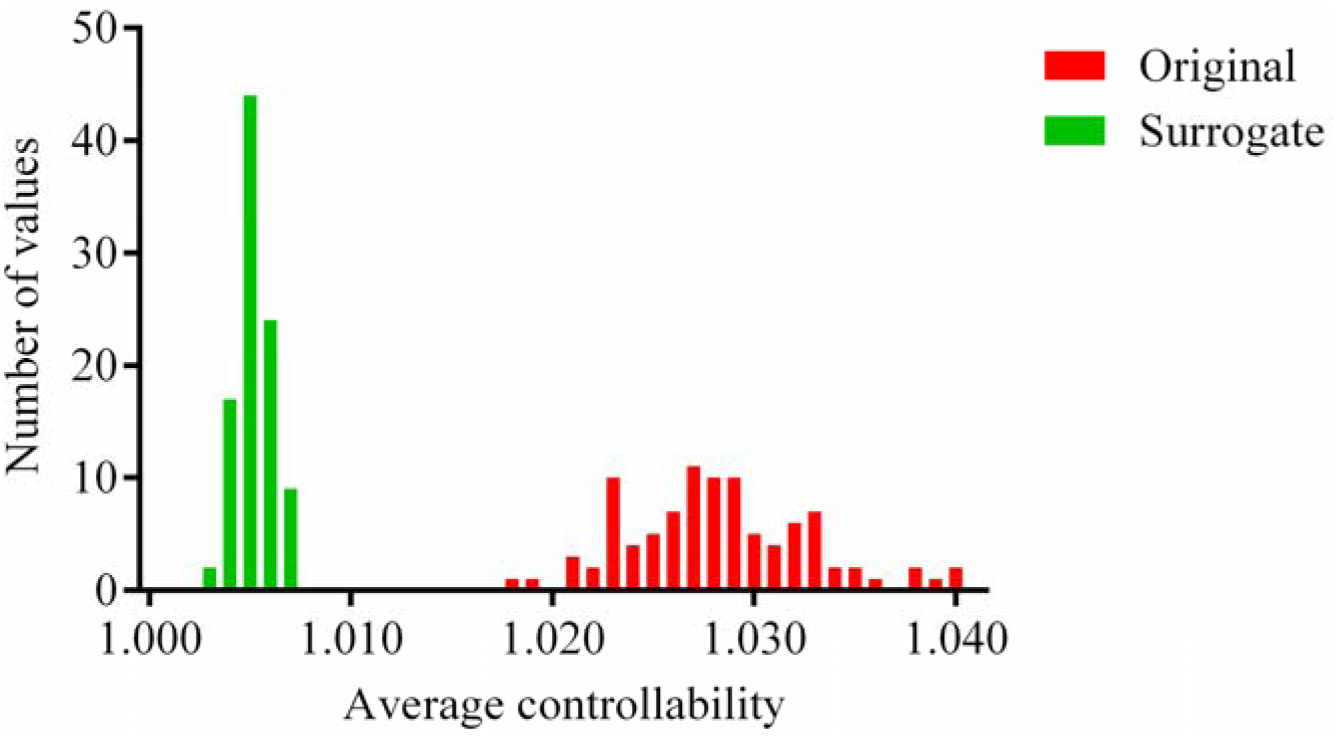
Distribution of average controllability values averaged across nodes in the original data (green) and in the random surrogate networks (red).

To quantify the contribution of topological factors to variability in controllability, stepwise linear regression was employed to model the putative contributions of various nodal measures (Table 2). As expected, nodal strength was the strongest predictor of average nodal controllability. However, other network features also make substantial contributions to controllability; the next most strongly predictive measure was subgraph centrality, followed by the clustering coefficient. Subgraph centrality is defined by the weighted sum of closed walks of different path lengths starting and ending at the nodal region (Estrada and Rodriguez-Velazquez, 2005). Clustering reflects the number of closed cycles with three edges. Average controllability, the impulse response to a stimulus, is hence magnified by positive feedback via closed walks and loops (Supplementary Figure 1).

**Table 2.** Stepwise linear regression of average controllability.

| Variable | Standardized<br>beta | Full correlation | Partial correlation |
| --- | --- | --- | --- |
| Node strength | 0.648** | 0.886 | 0.587 |
| Subgraph centrality | 0.297** | 0.608 | 0.600 |
| Clustering coefficient | 0.152** | 0.388 | 0.387 |
| Within-module z-score | 0.098* | 0.793 | 0.114 |

Adjusted $R^2 = 0.887$ .
Excluded variables: local efficiency, betweenness centrality, participation coefficient.
Significance levels: \* $p < 0.01$ , \*\* $p < 0.001$

### 3.2. Controllability in high-risk and bipolar disorder subjects

Having established that node strength, subgraph centrality, clustering coefficient, and within-module *z*-score contribute to network controllability in our data, we next studied between-group differences in controllability at the three levels of granularity.

#### 3.2.1. Node level

Six out of seven regions previously identified as having reduced node strength in HR and BD groups also show significant between-group controllability differences. Reductions in average controllability were observed in the left insula, left parahippocampal gyrus, left middle occipital gyrus, left superior frontal gyrus, right inferior frontal gyrus / pars triangularis, and right precentral gyrus (Supplementary Table 1).

#### 3.2.2. Subnetwork level

We next studied average controllability differences in four lateralized subnetworks (Fig. 2, Supplementary Table 2). Significant average controllability differences (*p*<0.05, corrected) were seen in subnetworks A and B, but not C or D (Fig. 5). Controllability of Network A, including the left superior/inferior frontal gyri, postcentral gyrus, insula, and pars triangularis (Fig. 1A), was reduced in BD compared to CN and HR (□^2^ =0.043, *p*<0.004 for BD vs. CN, *p*<0.024 for BD vs. HR). Average controllability of Network B, involving the right superior/middle/inferior frontal gyri, superior temporal pole, putamen, caudate, pars triangularis, and pars orbitalis (Fig. 1B) was reduced in HR subjects compared to CNs (*p*<0.008, □^2^ =0.037). Network C showed no significant group differences (*p*=0.544). Average controllability differences approached significance (p=0.059) between CN and the BD group in Network D.

**Figure 5.**
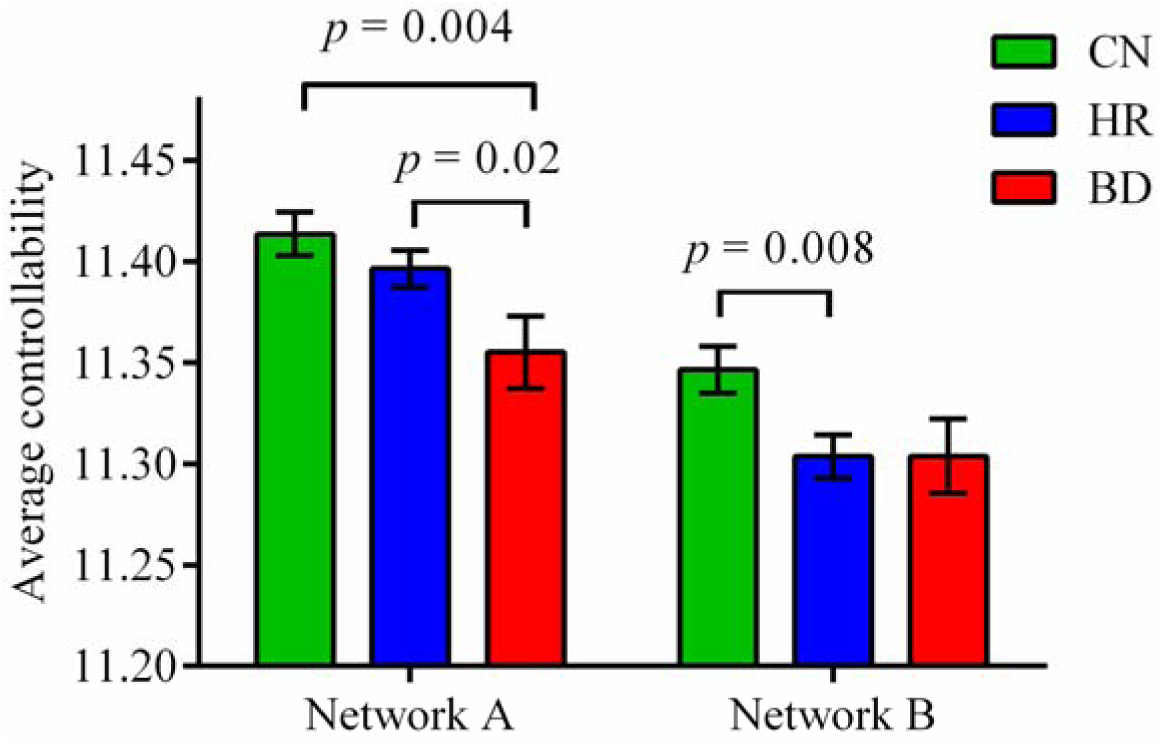
Mean average controllability for subnetworks with identified significant group differences. Error bars show standard error of the mean. CN, controls; HR, high-risk; BD, bipolar disorder.

These effects were very similar across a range of connection densities within the thresholded networks. The BD vs. HR contrast of Network A became nonsignificant for connection densities of > 15% (*p*=0.024 vs. 0.027, see Supplementary Table 3, 4 and 5).

Group differences in Networks A and B were reduced after strength-preserving randomization, highlighting the role of network topology (Table 2), consistent with (Kim *et al.*, 2018).

**Table 2.** Differences between cohort-mean controllability values for each group before and after strength-preserving randomization.

|  | Original | Surrogate |
| --- | --- | --- |
| Network A: CN-BD | 0.059 | 0.012 |
| Network A: HR-BD | 0.041 | 0.008 |
| Network B: CN-HR | 0.043 | 0.008 |

#### 3.2.3. Community level

We finally addressed average controllability at the scale of brain communities whose specialized functional roles have been previously documented (Yeo *et al.*, 2011). There were substantial differences in controllability values across these communities, with the limbic and frontoparietal modules showing the highest per-node average controllability (Fig. 6). These differences reflected, but were not strictly enforced by, differences in average node strength. For example, the limbic and frontoparietal communities switch relative positions of strength *versus* controllability. However, no group-wise differences in average controllability of these communities survived FDR correction.

**Figure 6.**
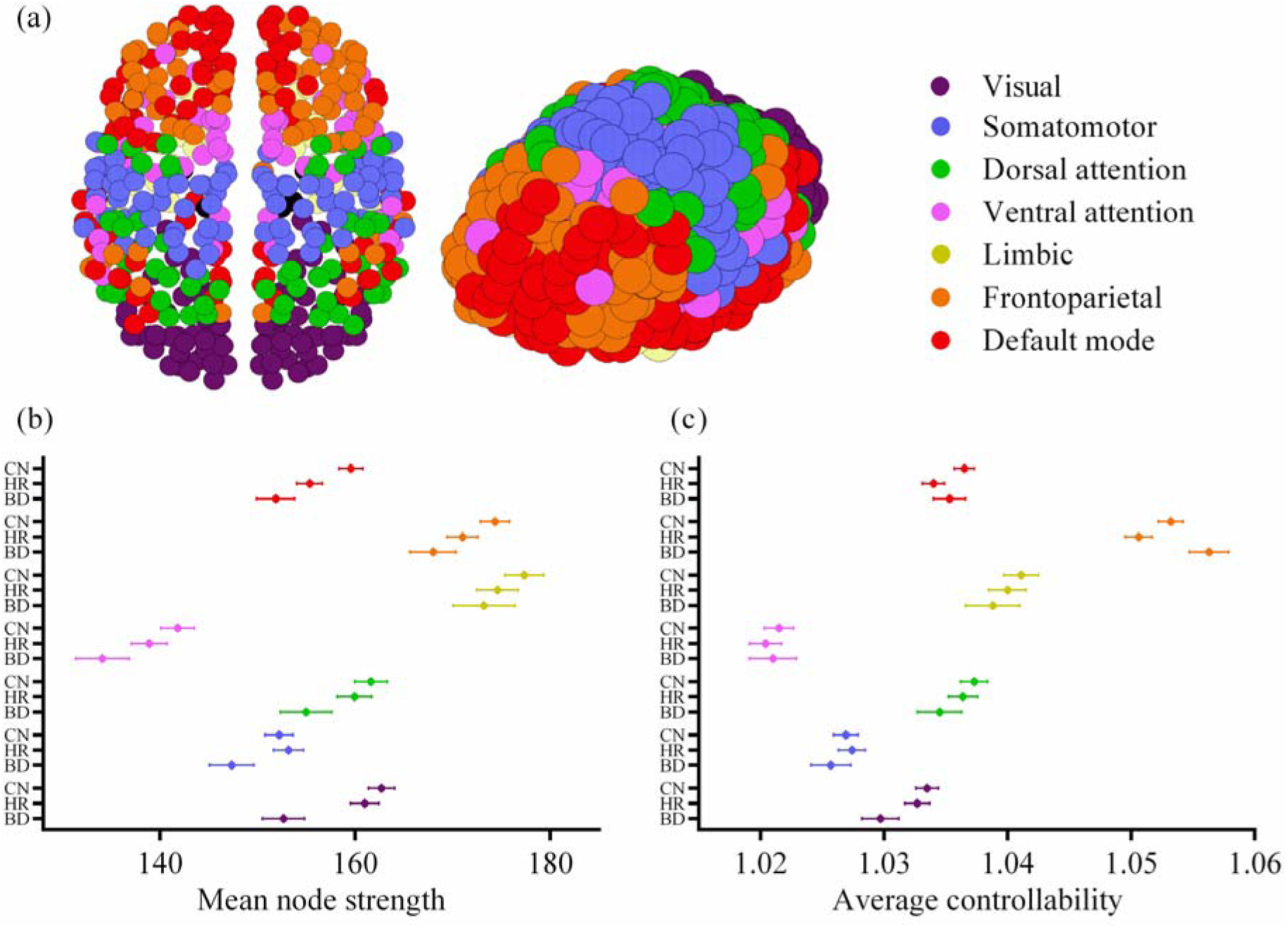
Average controllability of communities across the three groups. (a) Corresponding affiliation of each node to the functional communities derived from (Yeo *et al.*, 2011). (b) Mean node strength, and (c) average controllability (normalized by community size) in each group and each functional module. Errors bars show standard error of the mean. CN, controls; HR, high-risk; BD, bipolar disorder.

#### 3.2.4. Auxiliary analyses

We observed a significant relationship (after FDR-correction) between age and controllability in the CN group (*r*=0.291, *p*=0.004). However, for the group contrasts of interest, there were no significant age x group interactions for average controllability (Network A CN vs. BD *p*=0.07; Network A HR vs. BD *p*=0.09; Network B CN vs. HR *p*=0.93; Supplementary Figure 2).

The conversion from HR to BD corresponds to the occurrence of a manic or hypomanic episode, the incidence of which peaks in the third decade of life (Saunders and Goodwin, 2010). Future risk of conversion is therefore higher in our younger HR participants, reducing towards the upper age limit of the cohort. To study the network correlates of this effect, we stratified HR subjects by age, divided at the mean age of illness onset (< 22 or ≥ 22 years). The younger HR sub-cohort had reduced Network B controllability compared to the older group (*p*=0.02, Cohen’s *d* = 0.52). No difference was observed in Network A.

BD cleaves into two clinical subtypes, with psychotic symptoms occurring in the elevated (manic) phase of bipolar Type I but not during the (hypomanic) phase of those with bipolar Type II. Interestingly, participants with bipolar Type II had significantly weaker controllability for Network A than subjects with bipolar Type I (*p*=0.01, Cohen’s *d* = 0.88). No significant difference was seen for Network B.

The occurrence of a major depressive episode in HR individuals may represent an important early stage in the development of BD (Perich *et al.*, 2015). However, the average controllability for Networks A and B in HR individuals was not significantly modulated by a history of anxiety or major depression (Supplementary Table 9). Similarly, measures of illness severity in the BD group (illness duration and total number of mood episodes) were not significantly associated with the average controllability of the two subnetworks (Supplementary Table 10).

Even outside of frank mood episodes, patients with BD often describe emotional dysregulation. We thus also studied putative associations with the Cognitive Emotion Regulation Questionnaire scores (Garnefski and Kraaij, 2007) – a well validated self-rated measure of cognitive coping strategies. After FDR correction, two subscales of the CERQ were significantly different between groups, with BD subjects having low ‘positive refocusing’ and ‘high catastrophizing’ scores. Other subscales – self-blame, rumination, and positive appraisal – approached significance (Supplementary Table 11). Given the CERQ subscale scores are highly correlated, principal component analysis (PCA) was employed to reduce the dimensionality of the subscales to a smaller number of latent factors. The first principal component (i.e. factor), which captures the largest variance in the original CERQ subscales scores, was also significantly different between groups (*p*=0.0004). However, none of the subscales, nor the first principal component, was significantly correlated with controllability in Network A or in Network B within the HR or BD groups (*p*>0.327).

## 4. Discussion

Quantifying brain structural network disturbances in psychiatric conditions provides novel insights into their neurobiological correlates, yet lacks a direct explanatory link to the illness phenotype. We previously observed structural network changes in those with, or at high genetic risk of BD (Roberts *et al.*, 2016b). Here we used network control theory to address the theoretical gap between these structural network disturbances and the dysregulation of cognitive-emotional function that characterizes BD and those at HR. Specifically, we asked whether focal and distributed network disturbances observed in HR and BD subjects translated to impaired brain controllability as predicted by linear network control theory. Controllability deficits were seen in six out of seven key brain regions and also in two out of four subnetworks: a left-sided network involving the insula and postcentral gyrus in BD, and in a right-lateralized PFC-striatal subnetwork in HR. Therefore, group differences in network strength – seen in all four subnetworks - do not fully account for group differences in controllability. There were no group differences in controllability in seven widely studied functional communities, suggesting that the controllability deficits in BD originate focally rather than diffusely. We now consider the conceptual and clinical implications of these findings.

The average controllability quantifies the capacity of input to a subnetwork to manipulate the transition of broader brain states (Gu *et al.*, 2015). Given structural connectivity data, it predicts the network-wide impulse response to a stimulus applied to a node or subnetwork of interest, (Muldoon *et al.*, 2016). This interpretation is instructive, as signal spread depends on (i) amplification by positive feedback within the subnetwork, and (ii) links between the subnetwork and its surrounds (Kim *et al.*, 2018). We found that the top three predictors of average controllability in our data were node strength, subgraph centrality, and local clustering coefficient, highlighting the role of closed motifs such as triangles and larger loops in amplifying positive feedback, in addition to the role of strong connections in propagating that energy outwards. Topological factors such as these amplifying motifs were responsible for 32% of the between-subject variability in controllability. Removal of these topological factors by network randomization reduced between-subject and between-group differences. Hence the present work builds upon our previous observation of between-group subnetwork strength differences (Roberts *et al.*, 2016b) by showing how these, as well as other network features, translate into group differences in subnetwork average controllability.

Impaired controllability was seen in six brain regions and two lateralized networks. Although Network A (involving the left superior and inferior frontal gyri, insula, and postcentral gyrus) had reduced connectivity in HR compared to CN, it showed impaired controllability exclusively in the BD group. Despite the loss of power imposed by splitting our clinical BD group, Network A showed significantly weaker controllability in BD Type II than BD Type I, consistent with recent evidence of disparate underlying etiologies (Song *et al.*, 2018). Network A thus represents a putative subgroup biomarker, although this clearly requires replication in an independent data set. NBS identifies subnetworks by comparing mean edge weights across groups, but is insensitive to topological group differences which would also influence controllability. Such effects are also consistent with the loss of significant group differences with strength-preserving randomization. The regions and connections of Network A overlap with previous demonstrations of changes in fractional anisotropy for tracts involving the left superior frontal gyrus (Adler *et al.*, 2006), decreased effective connectivity between the left dorsolateral prefrontal cortex (PFC) and the left inferior frontal gyrus (Breakspear *et al.*, 2015), hypo-activation in the ventrolateral PFC during emotionally salient tasks (Chen *et al.*, 2011; Phillips and Swartz, 2014), and altered reward processing-related activity in the left striatum and left ventrolateral PFC (Bermpohl *et al.*, 2010; Nusslock *et al.*, 2012). Whereas the present study is only concerned with structural imaging data, these latter studies highlight corresponding changes in the activity and function of the same networks, possibly reflecting the consequences of network controllability. Altered connectivity between these frontal areas and the insula, a region involved in interoceptive processes (Critchley *et al.*, 2004), may mediate interplay between dysfunctional cognition and emotional homeostasis in BD. That is, impaired signal amplification from these circuits, arising from reduced controllability, could produce the loss of emotional control that characterizes BD.

We also observed reduced controllability in Network B (which encompassed the right frontal gyri, superior temporal pole, putamen, and caudate nucleus), in this case, for HR compared to CN. Existing research has also noted structural and functional changes in the right frontal white matter (Bruno *et al.*, 2008), right ventrolateral PFC (Hajek *et al.*, 2013), putamen and caudate (Haller *et al.*, 2011), and right amygdala (Torrisi *et al.*, 2013). PFC dysregulation has been a central theme in the psychiatric neuroimaging literature. This region is involved in executive function, cognition, planning, and reward. Attenuated controllability of this right-lateralized frontostriatal reward loop, seen in HR subjects alone, may reflect a marker of resilience or risk. Alternatively, psychotropic medications taken by the BD group may have altered structural connectivity (Dusi et al., 2015). However, both the HR and BD groups had the same mean controllability for this subnetwork (11.30 vs. 11.30) but only the CN-HR contrast was significant while CN-BD was not. The lack of significance for the BD contrast may reflect loss of power (associated with the smaller BD cohort) and/or increased inter-subject variance in the BD group. Also of interest, controllability of Network B was most strongly reduced in the younger sub-cohort of HR subjects, who have not yet passed the peak age of illness onset and are hence, on average, at higher future risk of BD compared to the older HR participants. Longitudinal follow-up will allow us to draw more definitive conclusions by seeing if individual differences in controllability predict later conversion to BD.

Although we also selected Networks C and D for further analyses based on our prior observation of group differences in intra-network connectivity (Roberts *et al.*, 2016b), we did not observe group differences in average controllability. Therefore, network strength differences alone are not sufficient to confer differences in controllability.

The different network controllability effects in the HR group compared to BD are intriguing. Controllability deficits that are specific to BD may reflect a purer form of illness risk and expression than those in HR, which rather reflect a mix of risk and resilience but not (yet) BD expression. Effects in BD alone may also be state markers of medication exposure or the consequences of recurrent mood episodes. The left-hemisphere disturbances seen in BD alone may reflect the left-lateralization of autonomic processes in fronto-temporal areas (Guo *et al.*, 2016). Moreover, the edges in Network A are more distributed across the cortex – potentially allowing propagation of fluctuations to cognitive, emotional, interoceptive, and somatosensory areas.

The objectives of the present study rested on the conjecture that diminished controllability in emotional and cognitive control networks would underlie the emotional dysregulation seen in BD and HR participants. Because the presence of a frank mood disturbance was an exclusion criteria for our study, we studied the Cognitive Emotion Regulation Questionnaire (CERQ) as a more nuanced proxy to this construct (in contrast to symptom scores). There were significant CERQ subscore differences between BD and CN, and to a lesser degree, between HR and CN. These parallel our group differences in controllability. However, controllability did not covary with CERQ scores within each group. It is possible that controllability captures the gross group differences in cognitive coping strategies between the groups, but not the more nuanced differences between individuals within these groups. A larger study may provide clarification as statistical power was limited by group size in HR and BD, as well as adjustment for multiple testing across both subnetworks and 9 subscales.

Several caveats of this study need to be considered. Interpreting subnetwork controllability as a surrogate for signal amplification assumes that external edges are preserved. Edges that are outside a subnetwork discovered through statistical testing may be weakened but with insufficient magnitude to be included in the final supra-threshold subnetwork. Therefore, average controllability should be conservatively interpreted as a combination of subnetwork outflows and internal signal amplification (Kim *et al.*, 2018). Selective analysis of controllability relies substantially, but not solely, on strength differences in the subnetworks identified previously (Roberts *et al.*, 2016b). However, we observed that there was a strong influence of topological features, which contribute to 32% of inter-subject variability. Subgraph centrality and the clustering coefficient were included as predictors in the regression model independent of their collinearity with node strength, demonstrating the influence of complex network motifs such as closed walks and loops. Therefore, the present analyses provide additional insights not provided through interrogation of edge strength alone.

Psychiatric comorbidity in the HR group is a potential confounder, with 8% of this group suffering recurrent major depressive episodes. However, controllability in the HR group did not co-vary with comorbid anxiety or depressive episodes, illness duration or total number of mood episodes. It is also important to note that prior depression or anxiety was not an exclusion factor for our control cohort, ensuring that a proportion of these participants also had some psychiatric comorbidity (and hence avoiding a “super-healthy” cohort). The BD cohort did not include any subjects with a current mood state of depression, hypomania, or mania. This, plus the use of clinically titrated medication, limits the interpretability of any putative associations with current symptom severity. Moreover, it seems unlikely that controllability, a measure derived from structural connectivity, would be altered in the short term by current mood state. Controllability hence appears to speak to trait rather than state differences between our three cohorts. Again, longitudinal follow-up will be useful in identifying the controllability precedents to the development of manic symptoms in the HR individuals who do transition to BD. Expansion to a larger, multi-disorder cohort – including patients with other psychotic disorders such as schizophrenia – would also allow testing for the specificity of controllability differences in the present subnetworks. In analogy to findings in structural connectivity, controllability differences in schizophrenia may be more distributed than those in bipolar disorder, possibly involving core structural networks, such the rich club (van den Heuvel *et al.*, 2013). However, any global measure of brain-wide controllability would be difficult to detect using the present algorithm, as the network is normalized to prevent unstable dynamics before estimating controllability values.

There were 5 families that included a member in the HR and in the BD groups. In the original discovery of structural network differences in this cohort, generalized estimating equations were used to account for the relatedness of the small number of family members in the cohort (Roberts et al., 2016b). However, practically we found use of this more complex statistical framework had a negligible effect on the reported group differences. Likewise, removing the family members from the analyses had no impact on the significance of the subnetwork effects. Therefore, in the present study, we employed a standard general linear model. Future work could focus on the subtle nuances of familial relatedness, although the need for a much larger cohort size would be mandate a consortia, data-sharing approach.

In addition, controllability is predicated upon the reduction of a complex nonlinear system to the simple linear model given in Equation (1). Several points pertain to this simplification. First, although fine-grained, fast dynamics bear the fingerprints of highly nonlinear dynamics such as multistability (Freyer *et al.*, 2011), linear models have been found to predict much of the slow fluctuations and time-averaged variance in empirical recordings (Honey *et al.*, 2009; Deco *et al.*, 2013). Second, complex nonlinear dynamics organize around simple underlying attractors, including fixed points (Cocchi *et al.*, 2017). Linear models provide a sufficient approximation for the behavior of dynamics in the neighborhood of such fixed points (Hirsch *et al.*, 2013). Linear theory, therefore, arguably provides a unique, albeit incomplete, account of brain network controllability. Future work could harness the potential of nonlinear control theory, although the challenges in doing so remain substantial (Slotine and Li, 1991; Cornelius *et al.*, 2013).

Finally, our interrogation of network controllability relied exclusively on predicted changes in network function from the analysis of structural connectivity data. Changes in task activation and resting state functional connectivity do occur in these same participants, overlapping partially with the networks studied here (Roberts *et al.*, 2013; Roberts *et al.*, 2016a). Correlating group differences in controllability with group differences in functional imaging would be instructive as both measures depend on the interplay of structural connectivity and local dynamics. However, it is crucial not to conflate the two. Functional connectivity reflects signal correlations between brain regions, as seen through the lens of hemodynamic responses to intrinsic fluctuations and sensory inputs. Changes in local and regional controllability may manifest as changes in distal connections, and may not necessarily produce one-to-one changes in regional functional connectivity (Tang *et al.*, 2018).

Quantifying structural network differences in high risk and psychiatric cohorts is an important step in developing novel diagnostic markers for risk and resilience. However, computational frameworks are required to understand the link between these illness correlates and the phenotype (Fornito *et al.*, 2015). In this vein, control theory can lend early insights into distinguishing between state markers of disease, trait markers, and even adaptive changes that may be harnessed in future clinical interventions. Network B, a putative frontostriatal reward loop, had attenuated controllability in the HR group alone. Longitudinal follow-up of the HR group is currently underway to assess whether network changes are protective adaptations or reflect trait risk to the disorder. The absence of controllability deficits in Networks C and D, despite edge strength deficits, suggests broader compensatory topological changes: This conjecture also suggests developmental processes that could be tested through longitudinal follow-up. Alternatively, targeted non-invasive stimulation of these brain regions in healthy subjects represents another means of testing such predictions. Non-invasive stimulation of weakened networks holds theoretical promise to mitigate disease transition in high-risk individuals, or perhaps even restore cognitive-emotional control in BD patients.

## Funding and acknowledgements

We would like to thank the participants and staff of the Kids and Sibs study for their invaluable contribution towards the project. This study was funded by the Australian National Medical and Health Research Council (118153, 10371296, 1095227), the Lansdowne Foundation and Brother’s Reid. DSB also acknowledges support from the Alfred P. Sloan Foundation, the John D. and Catherine T. MacArthur Foundation, and the ISI Foundation. The content is solely the responsibility of the authors and does not necessarily represent the official views of any of the funding agencies.

## Abbreviations

CN: control;
HR: high-risk;
BD: bipolar disorder;
MRI: magnetic resonance imaging;
fMRI: functional MRI;
dMRI: diffusion MRI.

