## Supplementary Materials for "Fronto-limbic dysconnectivity leads to impaired brain network controllability in young people with bipolar disorder and those at high genetic risk"

### Supplementary Material

#### Normalization of networks

Controllability analyses are predicated upon linear stability of all individual subject networks, which was ensured by dividing all edge weights by a number 10% greater than the largest eigenvalue among all subjects. We chose to rescale all subjects by a common number rather than normalize each subject separately to preserve individual and group differences in mean weighted connectivity. We chose 10% to prevent unstable dynamics in the most strongly-connected (least stable) subject. Results were consistent across greater degrees of network stabilization (Supplementary Table 6).

#### Network controllability

The eigenvector states v(t) of each network are derived from this equation by linear algebra. These are patterns of activation satisfying the relation x(t+1)=Av(t)=μv(t), where the eigenvalue μ is a constant. Once an eigenvector state is initiated, the relative activation of different nodes within the network will remain unchanged. However, the magnitude of v(t) will be amplified (μ>1) or damped (μ<1) with time.

The energy required to reach a target state is obtained by summing energy input over time. Let the energy of the control input be $E\left( u \right)= \sum_{t=0}^{N-1} \left\| u(t) \right\|^{2}$, the sum of the energy applied to the input nodes. The minimum energy control input that takes the network from the zero state $x\left( 0 \right)=0$ to the target state $x\left( N \right)=x_{f}$ is,

u^*^(t) = B^T^(A^T^)^N-t-1^W_K_^-1^ x_f_,	(2)

where $W=\sum_{t=0}^{\infty} A^{t}BB^{T}A^{t}$ refers to the controllability Gramian.

One can show that$E(u^{*})\leq{1/\mu}_{min}$, where $\mu_{min}$ is the smallest eigenvalue of $W$. Therefore, µ_min_ sets the energy requirement of the most difficult-to-reach state.

Average controllability for a set of control nodes is the average energy needed to steer the system to any target state in finite time. This is directly proportional to Trace${(W}^{-1})$. We use Trace$(W)$, the mean impulse response energy, as the measure of average controllability because Trace$(W)$ and Trace${(W}^{-1})$ are highly correlated due to a relation of inverse proportionality, and Trace${(W}^{-1})$ is ill-defined. When the control set is a single node *i*, $B$ is just the input vector $e_{i}$to the $i$^th^ control node, and Trace$(W)$, the average controllability of node *i*, simplifies to,

$\text{Trace}\left( W \right)=\sum_{j} \frac{v_{\mathrm{ij}}}{1-{\mu_{j}}^{2}},$	(3)

where $i$ is a node, $v_{\mathrm{ij}}$ is the i^th^ component of the *j*^th^ eigenvector, and $\mu_{j}$ is the *j*^th^ eigenvalue. The variable $v_{\mathrm{ij}}$ can be considered as the influence of the $i$^th^ node on the $j$^th^ eigenmode.

#### Between-subjects and between-nodes variability in controllability

Controllability for each of the 512 nodes in each of the 96 control subjects fits in a 512 $\times$ 96 matrix (matrix *A*). Strength-preserving randomization generates an ensemble of surrogate networks, one from each of the 96 controls. The controllability values for each node in each of these surrogate networks fit in a new 512 $\times$ 96 matrix (matrix *B*). Spearman’s rank correlation between the *n*^th^ row of this matrix gives the correlation coefficient for the *n*^th^ node across all subjects. Averaging such values across all the 512 nodes yields the *between-subjects variability*. Conversely, the Spearman’s rank correlation between the *n*^th^ column of this matrix gives the correlation coefficient for the *n*^th^ subject across all nodes. Averaging such values across all the 96 control subjects yields the *between-nodes variability*.

#### Relationship between node strength and average controllability

Let *x* and *y* be canonical vectors corresponding to two nodes, such that node *x* has greater weighted degree than *y*. Then *x* has a greater impulse response ||*Ax*|| > ||*Ay*||. Hence,

$\left\| V^{T} \right\|\left\| \mathrm{Ax} \right\|>\left\| V^{T} \right\|\left\| \mathrm{Ay} \right\|$.	(1)

It then follows that,

$\left\| V^{T}\mathrm{Ax} \right\|>\left\| V^{T}\mathrm{Ay} \right\|.$	(2)

Let D be the diagonal matrix of eigenvalues. $Then D=V^{-1}AV and likewise, DV^{-1}=V^{-1}A$, where the vector V is orthonormal because A is symmetric,

$DV^{T}=V^{T}A.$	(3)

Substituting (3) into (1) yields,

$\left\| DV^{T}x \right\|>\left\| DV^{T}y \right\|.$	(4)

Expanding DV^T^ yields,

$\sum_{j} \mu_{j}{v_{j}}^{T}x>\sum_{j} \mu_{j}{v_{j}}^{T}y$	(5)

where $\mu_{j}$is the eigenvalue of column eigenvector $v_{j}$. Expanding the dot product,

$\sum_{j} \mu_{j}\left\| v_{j} \right\|\left\| x \right\|\cos\left( v_{j},x \right)>\sum_{j} \mu_{j}\left\| v_{j} \right\|\left\| y \right\|\cos\left( v_{j},y \right)$.	(6)

Hence we find that,

$\sum_{j} \mu_{j}\cos\left( v_{j},x \right)>\sum_{j} \mu_{j}\cos\left( v_{j},y \right) as x,y and v_{j} are unit vectors$	(7)

Therefore, higher-degree nodes (such as node *x*) are disproportionately larger components of high-eigenvalue eigenvectors, enhancing their average controllability $\sum_{j} \frac{v_{\mathrm{xj}}}{1-{\mu_{j}}^{2}}$.

#### Supplementary Data

#### Node controllability

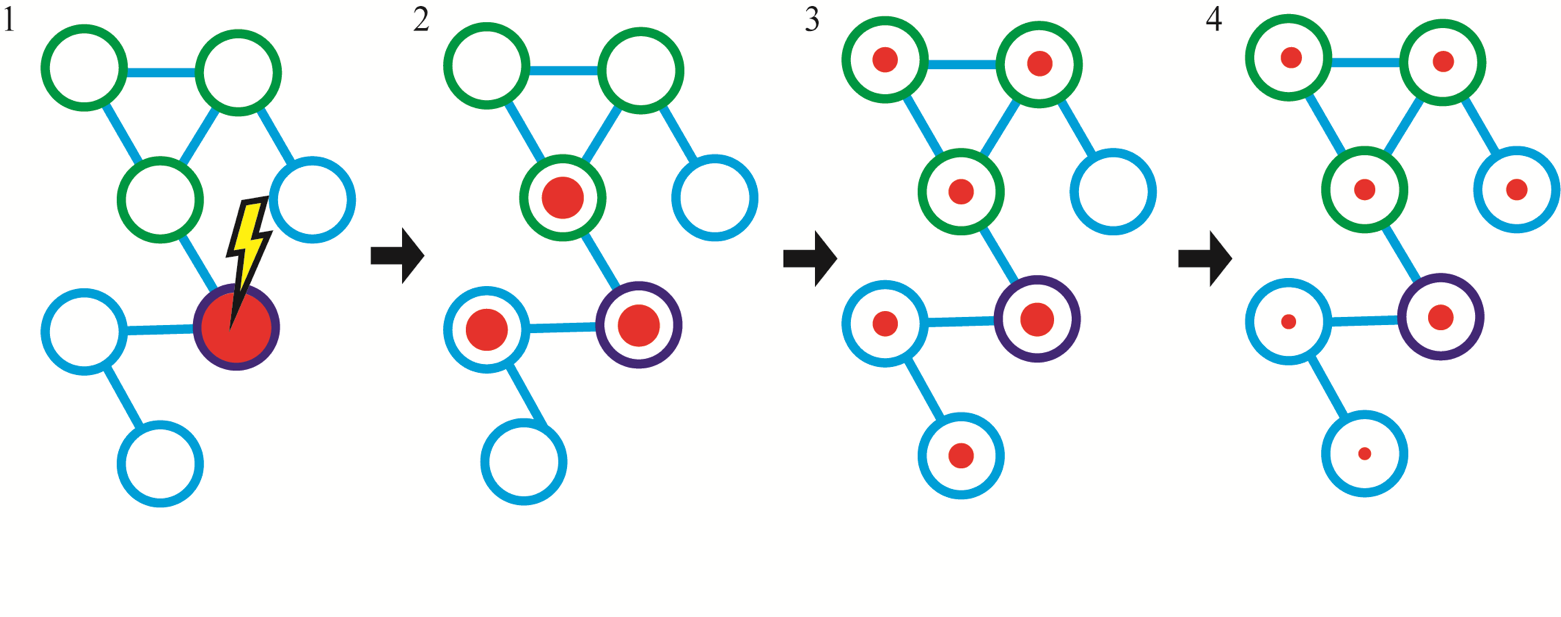

**Supplementary Figure 1.** Signal spread from one starting node (purple) in a brain network. The area of the red circle indicates the magnitude of activation. Positive feedback in closed loops (green nodes) slows signal decay.

**Supplementary Table 1.** Average controllability for nodes in CN, HR, and BD groups with significant between-group differences as identified by network-based statistic in the previous investigation (Roberts *et al.*, 2016b)

| Region | Node | CN | HR | BD | *p* | CN>HR | CN>BD | HR>BD |
| --- | --- | --- | --- | --- | --- | --- | --- | --- |
| Left Hippocampus | 83 | 1.014 | 1.011 | 1.010 | 0.066 | N/A | N/A | N/A |
| Left Insula | 197 | 1.021 | 1.019 | 1.015 | 0.006* | - | 0.002* | 0.014* |
| Left Parahippocampal Gyrus | 198 | 1.023 | 1.018 | 1.020 | 0.013* | 0.001* | - | - |
| Left Middle Occipital | 202 | 1.046 | 1.041 | 1.038 | 0.037* | 0.028* | 0.014* | - |
| Left Superior Frontal Gyrus | 211 | 1.042 | 1.032 | 1.030 | <0.001* | <0.001* | 0.001* | - |
| Right Inferior Frontal Gyrus / Pars Triangularis | 504 | 1.038 | 1.031 | 1.034 | 0.038* | 0.005* | - | - |
| Right Precentral Gyrus | 511 | 1.025 | 1.022 | 1.018 | 0.006* | - | 0.001* | 0.027* |

CN, Controls; HR, High-risk; BD, Bipolar Disorder

**p*<0.05 with Benjamini-Hochberg FDR correction.

#### Subnetwork controllability at main threshold and alternate graph sparsities.

**Supplementary Table 2.** Average controllability for subnetworks with groupwise differences in structural connectivity as identified in the previous investigation (Roberts *et al.*, 2016b). Network density is 10%. All networks normalized by 10% greater than the largest eigenvalue among all subjects.

| Subnetwork |  | CN | HR | BD | *p* | ƞ^2^ | CN vs HR^1^ | CN vs BD | HR vs BD |
| --- | --- | --- | --- | --- | --- | --- | --- | --- | --- |
| A |  | 11.414 | 11.397 | 11.355 | 0.009* | 0.043 | 0.22 | 0.004* | 0.024* |
| B |  | 11.347 | 11.304 | 11.304 | 0.017* | 0.037 | 0.008* | 0.052 | 1 |
| C |  | 10.132 | 10.131 | 10.123 | 0.544 | N/A | N/A | N/A | N/A |
| D |  | 6.073 | 6.067 | 6.061 | 0.059 | N/A | N/A | N/A | N/A |

CN, Controls; HR, High-risk; BD, Bipolar Disorder

^1^ Two-tailed *t*-test for each group contrast

**p*<0.05 with Benjamini-Hochberg FDR correction

**Supplementary Table 3.** Average controllability for 5% network density.

| Subnetwork |  | CN | HR | BD | *p* | CN vs HR^1^ | CN vs BD | HR vs BD |
| --- | --- | --- | --- | --- | --- | --- | --- | --- |
| A |  | 11.406 | 11.390 | 11.349 | 0.007* | 0.251 | 0.004* | 0.019* |
| B |  | 11.342 | 11.300 | 11.300 | 0.017* | 0.008* | 0.052 | 0.995 |
| C |  | 10.131 | 10.13 | 10.121 | 0.540 | N/A | N/A | N/A |
| D |  | 6.071 | 6.068 | 6.06 | 0.058 | N/A | N/A | N/A |

**Supplementary Table 4.** Average controllability for 15% network density.

| Subnetwork |  | CN | HR | BD | *p* | CN vs HR^1^ | CN vs BD | HR vs BD |
| --- | --- | --- | --- | --- | --- | --- | --- | --- |
| A |  | 11.415 | 11.398 | 11.357 | 0.010* | 0.225 | 0.005* | 0.027 |
| B |  | 11.347 | 11.304 | 11.304 | 0.017* | 0.008* | 0.052 | 0.992 |
| C |  | 10.133 | 10.132 | 10.123 | 0.546 | N/A | N/A | N/A |
| D |  | 6.073 | 6.069 | 6.061 | 0.059 | N/A | N/A | N/A |

**Supplementary Table 5.** Average controllability for full network density.

| Subnetwork |  | CN | HR | BD | *p* | CN vs HR^1^ | CN vs BD | HR vs BD |
| --- | --- | --- | --- | --- | --- | --- | --- | --- |
| A |  | 11.416 | 11.399 | 11.358 | 0.010* | 0.221 | 0.005* | 0.029 |
| B |  | 11.348 | 11.305 | 11.304 | 0.017* | 0.008* | 0.052 | 0.991 |
| C |  | 10.132 | 10.132 | 10.123 | 0.547 | N/A | N/A | N/A |
| D |  | 6.073 | 6.069 | 6.061 | 0.059 | N/A | N/A | N/A |

**Supplementary Table 6.** Average controllability for 10% network density, using a normalization factor that is 50% greater than the largest eigenvalue among all subjects.

| Subnetwork |  | CN | HR | BD | *p* | CN vs HR^1^ | CN vs BD | HR vs BD |
| --- | --- | --- | --- | --- | --- | --- | --- | --- |
| A |  | 11.196 | 11.189 | 11.169 | 0.004* | 0.312 | 0.002* | 0.009* |
| B |  | 11.169 | 11.149 | 11.149 | 0.011* | 0.006* | 0.039 | 0.977 |
| C |  | 10.067 | 10.067 | 10.063 | 0.494 | N/A | N/A | N/A |
| D |  | 6.037 | 6.035 | 6.031 | 0.050 | N/A | N/A | N/A |

#### Subnetwork node strength differences

**Supplementary Table 7.** Sum of node strengths for nodes in subnetworks with groupwise differences in structural connectivity.

| Subnetwork |  | CN | HR | BD | *p* | CN vs HR^1^ | CN vs BD | HR vs BD |
| --- | --- | --- | --- | --- | --- | --- | --- | --- |
| A |  | 2132 | 2056 | 1954 | <0.001* | 0.019* | 0.001* | 0.010* |
| B |  | 1958 | 1815 | 1806 | <0.001* | < 0.001* | 0.005* | 0.863 |
| C |  | 1308 | 1298 | 1247 | < 0.001* | 0.689 | 0.071 | 0.193 |
| D |  | 805 | 781 | 736 | <0.001* | 0.190 | 0.003* | 0.065 |

CN, Controls; HR, High-risk; BD, Bipolar Disorder

^1^ Two-tailed *t*-test for each group contrast

**p*<0.05 with Benjamini-Hochberg FDR correction

#### Subnetwork internal strength differences

**Supplementary Table 8.** Mean connections strengths for links in subnetworks with groupwise differences in structural connectivity.

| Subnetwork |  | CN | HR | BD | *p* | CN vs HR^1^ | CN vs BD | HR vs BD |
| --- | --- | --- | --- | --- | --- | --- | --- | --- |
| A |  | 0.392 | 0.230 | 0.318 | <0.001* | <0.001* | 0.072 | 0.005* |
| B |  | 2.827 | 1.948 | 2.132 | <0.001* | <0.001* | <0.001* | 0.216 |
| C |  | 0.387 | 0.617 | 0.471 | <0.001* | <0.001* | 0.033* | 0.024* |
| D |  | 2.473 | 2.271 | 1.735 | <0.001* | 0.096 | <0.001* | <0.001* |

CN, Controls; HR, High-risk; BD, Bipolar Disorder

^1^ Two-tailed *t*-test for each group contrast

**p*<0.05 with Benjamini-Hochberg FDR correction

#### Effects of comorbid anxiety/depression and age

**Supplementary Table 9.** Subnetwork average controllability for HR subjects stratified by anxiety experience and major depressive episode (MDE). There were no significant differences after FDR correction.

|  | Absent | Present | *p* |
| --- | --- | --- | --- |
| Subnetwork A: Anxiety | 11.388 | 11.443 | 0.022 |
| Subnetwork B: Anxiety | 11.296 | 11.345 | 0.091 |
| Subnetwork A: MDE | 11.398 | 11.393 | 0.835 |
| Subnetwork B: MDE | 11.312 | 11.279 | 0.182 |

**Supplementary Table 10.** Pearson correlation between subnetwork average controllability and measures of illness severity in BD subjects. N=37 for illness duration. N=35 for number of depressive episodes. There were no significant differences after FDR correction.

|  | R | *p* |
| --- | --- | --- |
| Subnetwork A: illness duration | 0.0057 | 0.973 |
| Subnetwork B: illness duration | -0.996 | 0.558 |
| Subnetwork A: number of depressive episodes | 0.1059 | 0.545 |
| Subnetwork B: number of depressive episodes | 0.1599 | 0.359 |

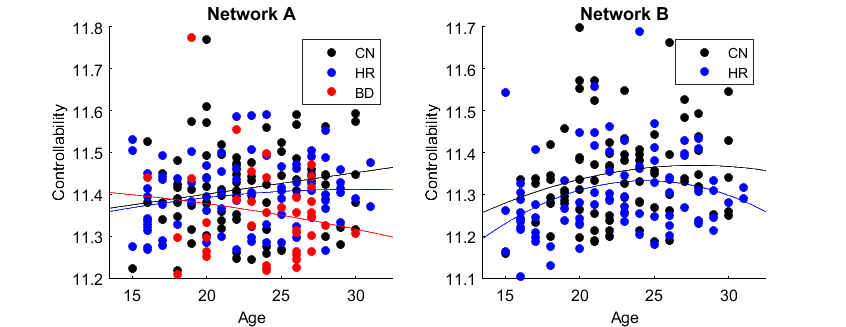

**Supplementary Figure 2.** Relationship between age and subnetwork average controllability for each significant group contrasts in previously identified subnetworks.

**Supplementary Table 11.** Cognitive Emotional Regulation Questionnaire (CERQ) scores

| Subscale |  | CN | HR | BD | *p* |
| --- | --- | --- | --- | --- | --- |
| Self-blame |  | 10.827 | 10.795 | 12.310 | 0.065 |
| Acceptance |  | 13.8 | 14.247 | 13.828 | 0.548 |
| Rumination |  | 12.653 | 12.521 | 14.276 | 0.038 |
| Positive refocusing |  | 12.227 | 11.795 | 9.862 | 0.007* |
| Refocus on planning |  | 15.2 | 14.904 | 13.517 | 0.045 |
| Positive reappraisal |  | 15.4 | 14.836 | 13.482 | 0.026 |
| Putting into perspective |  | 14.653 | 14.986 | 13.000 | 0.023 |
| Catastrophizing |  | 8.227 | 8.480 | 10.724 | 0.004* |
| Other-blame |  | 8.827 | 9.123 | 8.724 | 0.755 |
| First principal component |  | 16.137 | 15.667 | 11.369 | <0.001* |

CN, Controls; HR, High-risk; BD, Bipolar Disorder

**p*<0.05 with Benjamini-Hochberg FDR correction
